# Low-dimensional neural manifolds for the control of constrained and unconstrained movements

**DOI:** 10.1101/2023.05.25.542264

**Authors:** Ege Altan, Xuan Ma, Lee E. Miller, Eric J. Perreault, Sara A. Solla

## Abstract

Across many brain areas, neural population activity appears to be constrained to a low-dimensional manifold within a neural state space of considerably higher dimension. Recent studies of the primary motor cortex (M1) suggest that the activity within the low-dimensional manifold, rather than the activity of individual neurons, underlies the computations required for planning and executing movements. To date, these studies have been limited to data obtained in constrained laboratory settings where monkeys executed repeated, stereotyped tasks. An open question is whether the observed low dimensionality of the neural manifolds is due to these constraints; the dimensionality of M1 activity during the execution of more natural and unconstrained movements, like walking and picking food, remains unknown. We have now found similarly low-dimensional manifolds associated with various unconstrained natural behaviors, with dimensionality only slightly higher than those associated with constrained laboratory behaviors. To quantify the extent to which these low-dimensional manifolds carry task-relevant information, we built task-specific linear decoders that predicted EMG activity from M1 manifold activity. In both settings, decoding performance based on activity within the estimated low-dimensional manifold was the same as decoding performance based on the activity of all recorded neurons. These results establish functional links between task-specific manifolds and motor behaviors, and highlight that both constrained and unconstrained behaviors are associated with low-dimensional M1 manifolds.

## INTRODUCTION

The number of neurons involved in planning and executing a motor behavior, no matter how simple or complex, far exceeds the number of variables relevant to behavior. This mismatch makes the signals across neurons highly redundant and raises the question of how the activity of a population of neurons represents the relatively few variables relevant to behavior^1–3^. Contemporary studies across many brain areas report that the activity of populations of neurons is constrained to low-dimensional subregions of the neural space known as neural manifolds^2,4–20^: the computations required for planning and executing behaviors appear to be carried out through the patterns of neural activity within the manifolds rather than through the independent activity of individual neurons^8,13,14,17,21,22^.

Despite widespread use of the term “low dimensionality”, its precise meaning and significance remains unclear. Specifically, it raises questions such as: What qualifies as low dimensional? How does the behavioral complexity affect the dimensionality of the neural manifold? Should the nonlinearity in the neural manifold be considered when estimating its dimensionality? Recent definitions from Jazayeri and Ostojic’s comprehensive review aim to clarify this ambiguity with a focus on the neural computational principles associated with both *intrinsic* and *embedding dimensionality* in neural recordings^23^. A simple example illustrates the distinction between intrinsic and embedding dimensionalities (Jazayeri and Ostojic 2021). A ring has an intrinsic dimension of 1. If it is flat, it lies in an (*x*_1_, *x*_2_) plane, and its embedding dimension is 2. If the ring is made of a flexible material, it can be locally pinched away from the (*x*_1_, *x*_2_) plane. A full description of the ring now requires an additional direction *x*_3_; its embedding dimension is 3. If the ring existed within a 100-dimensional space, there are still 97 directions along which the ring could be locally pinched away from the (*x*_1_, *x*_2_) plane. Each such pinch involving a new direction increases the embedding dimension by one without affecting the intrinsic dimension. This simple example illustrates the intuition that the discrepancy between embedding and intrinsic dimensions is a proxy for the degree of nonlinearity of the manifold^24^.

The geometrical distinction between intrinsic and embedding dimensions leads to recent hypotheses about their distinct functionalities: the intrinsic dimensionality quantifies the number of independent latent variables needed to describe the collective activity of a population of neurons, while the embedding dimensionality is related to how collective information is processed and relayed downstream (Jazayeri and Ostojic 2021). For instance, recent theoretical studies posit that the low embedding dimensionality of the relevant manifolds allows for accurate brain-to-behavior maps^1,25^. A low embedding dimensionality implies that the relevant population dynamics can be captured by sampling the activity of a relatively small number of neurons. This observation has two implications. At a conceptual level involving our understanding of information processing in the brain, a low embedding dimension may facilitate the downstream readout of neural activity across brain areas; at a practical level involving our efforts to monitor and interpret neural population activity, a low embedding dimension allows the relevant population activity to be captured by the current recording technologies^26^.

The comparative analysis of intrinsic and embedding dimensionality is thus a critical concept to 1) quantify the redundancy in neural representations, 2) identify the number of latent components of collective activity in the neural population, 3) investigate the relation between these latent variables and behavioral task variables, sensory inputs, context, planning, and future expectations, and 4) quantify the extent to which behaviors can be decoded using current neural recording technologies.

In this study, we focus on primary motor cortical (M1) neural manifolds, which have been previously found to be low dimensional^2,9,24,27,28^. However, these earlier studies were conducted in laboratory settings where highly trained monkeys performed simple, constrained reach and grasp tasks. In contrast, natural behaviors outside the laboratory do not share these constraints. It remains unclear whether the observed low dimensionality of M1 manifolds is simply a byproduct of the constraints associated with laboratory tasks or if it reflects an intrinsic property of population dynamics in M1^1^. If the M1 manifolds corresponding to unconstrained behaviors were found to be similarly low dimensional, it would suggest that low dimensionality is a general organizational principle about neural population activity in M1. In this view, the relatively few latent variables that characterize a manifold would be sufficient for characterizing the cortical control of movement. These population signals are to be read out by the downstream neural circuitry that ultimately causes movement.

In intracortical Brain-Computer Interface (iBCI) applications, we operate under hardware limitations that vastly undersample the population of motor cortical neurons contributing to a specific behavior. When M1 population activity is constrained to a low-dimensional manifold, as is the case for constrained laboratory tasks, this limitation is overcome when decoding M1 neural activity within iBCIs. The question that we address here is whether the confinement of M1 population dynamics to low-dimensional manifolds also applies to unconstrained, naturalistic behaviors for which the development of iBCIs is highly desirable.

Our primary objective in this work was to characterize the intrinsic and embedding dimensionality of M1 manifolds corresponding to unconstrained behaviors such as grasping small treats while standing in the cage and quadrupedal locomotion over perch bars, and to compare them to the intrinsic and embedding dimensionalities of manifolds corresponding to constrained behaviors in the laboratory. In this analysis, we first applied denoising algorithms to the neural signals; this mitigates the overestimation effects of noise on the dimensionality estimates^24,29,30^. Following denoising, we computed both intrinsic and embedding dimensionalities of the neural activity. We found that the intrinsic and embedding dimensionality of neural manifolds were slightly higher in unconstrained settings, but still extremely low.

Our secondary objective was to characterize the geometry of the manifolds associated with unconstrained behaviors. To this end, we assessed the nonlinearity of the neural representations in both constrained and unconstrained settings. While we found evidence that the low-dimensional manifolds associated with unconstrained behaviors were nonlinear, most latent dynamics involved exploring nearly linear regions within the neural manifolds.

Our final objective was to investigate whether the low-dimensional M1 manifolds carry sufficient information about behavior to decode simultaneously recorded electromyograms (EMGs). In both laboratory and cage settings, we demonstrated that EMGs could be decoded from the level of activation of relatively few latent variables within neural manifolds, and that the performance of these decoders was as good as that obtained when decoding EMGs from the activity of all recorded neurons.

Our study illustrates that the low dimensionality of primary motor cortical manifolds is not exclusive to constrained laboratory tasks but is also present in unconstrained motor behaviors within a cage environment. Although the manifolds associated with unconstrained tasks have slightly higher intrinsic and embedding dimensionalities than those associated with constrained tasks, their existence indicates that the low dimensionality of M1 manifolds is characteristic of the population dynamics of M1 rather than a mere consequence of laboratory constraints. Our study advances our understanding of the computational strategies implemented to achieve M1’s role in processing and representing muscle-related information. In addition, we expect these findings to facilitate the extension of neural prosthetics and brain-machine interfaces to unconstrained settings.

## METHODS

### Recordings, tasks, and data preprocessing

We trained two 9-10 kg monkeys (Macaca mulatta) to perform tasks in two different environments. The first is defined as the “in-lab” environment, where the monkeys were seated on a standard primate chair and trained to perform a grasping task (**Fig 1a**). The task required them to reach and grasp a force-instrumented device located 30 cm in front of their shoulder using either left hand (monkey G) or right hand (monkey P). The shape of the device during an experimental session determined the type of grasping: a cylinder for power grasps with the palm and the fingers, a small rectangular cuboid for key grasps with the thumb and the edge of the index finger, and a small rectangular cuboid recessed within a thin slot for precision grasps with the tips of the thumb and the index finger. A pair of force sensitive resistors (FSRs) were attached on the sides of the devices to measure the grasping forces the monkeys applied. A monitor was placed above the device to display such forces with a cursor; the position of the cursor along the vertical and horizontal axes was determined by the sum and the difference of the FSR outputs, respectively. In each trial, the monkeys were initially required to keep the hand resting on a touch pad for a random time (0.5-1.0 s). A successful holding triggered the onset of one of three possible rectangular targets on the screen and an auditory go cue. The monkey was required to place the cursor into the target and to hold it there for 0.6 s by increasing and maintaining the grasping force applied on the device.

**Fig 1:**
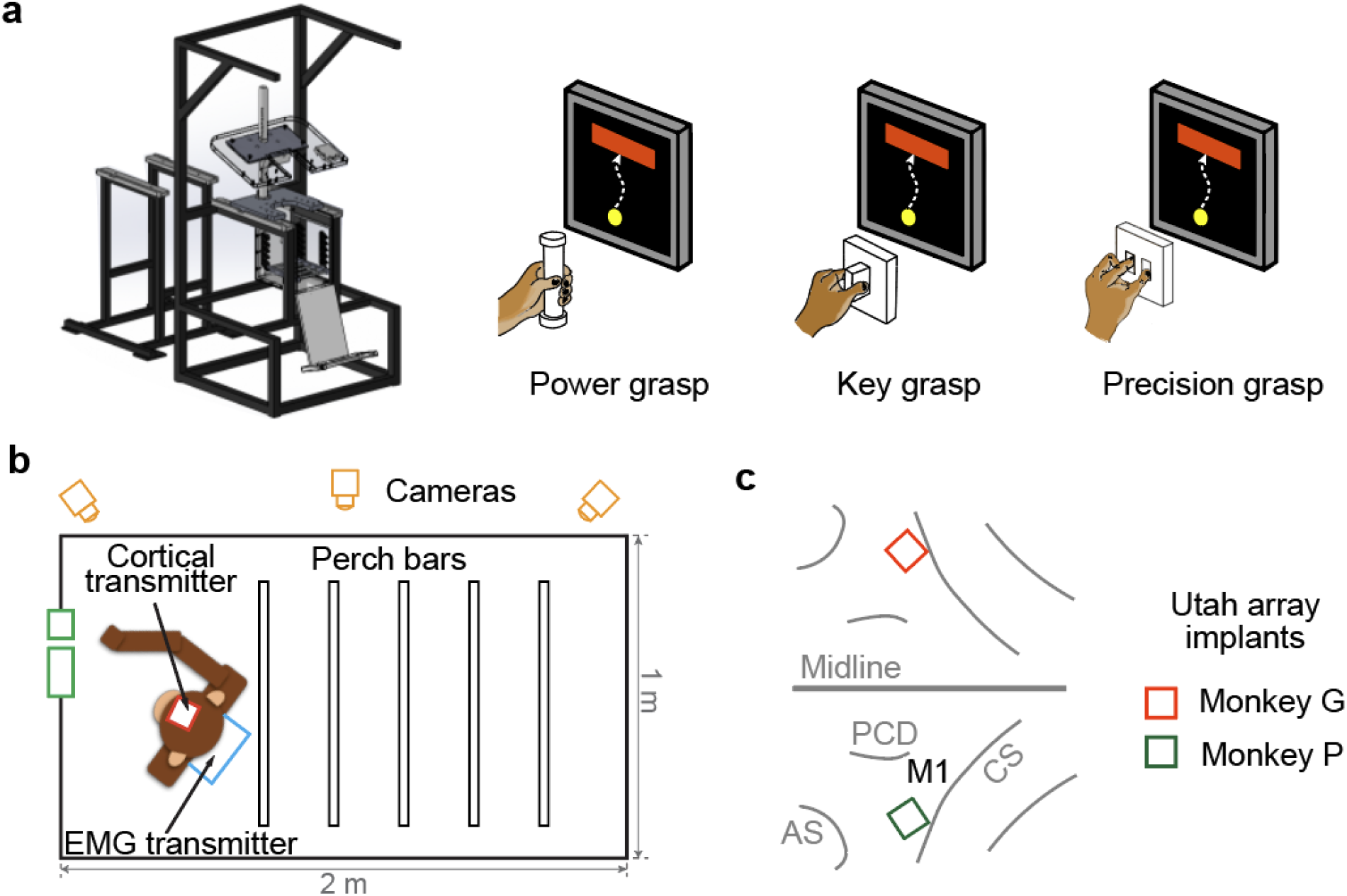
Tasks and recordings. a) Primate chair in which the monkeys performed a series of power, key, and precision grasp tasks in the constrained laboratory environment. b) The unconstrained cage environment where the monkeys moved freely. In the cage, monkeys did quadrupedal locomotion by grasping and walking over the perch bars (bar walk task). The monkeys also received treats from the experimenters (treat task). Monkeys were monitored using multiple synchronized cameras from which the individual behavior types were manually segmented, allowing for the corresponding segmentation of neural and EMG data. c) 96-electrode Utah arrays were implanted in the hand area of the primary motor (M1) cortex of each monkey (monkey G: right hemisphere, monkey P: left hemisphere).

The second environment is defined as the “in-cage” environment, where the monkeys were placed inside a 2×1×1 m plastic cage (**Fig 1b**). There were five bars spanning the width of the cage mounted 10 cm above the floor; the monkey grasped these bars while walking back and forth the length of the cage. The monkey typically would do a series of power grasps on these perches as he moved. We called this behavior *bar walk*. There are small holes on the door of the cage, through which experimenters could present small food pellets as treats to the monkeys inside; the monkeys typically used a precision grasp with the thumb and the index finger to take them. We called this behavior *treat grasp*. We identified single in-cage bar walk and treat grasp segments from the continuous recordings based on synchronized video recordings, and defined those segments as “successful trials”.

We implanted a 96-electrode array with 1.5 mm shaft length (Blackrock Microsystems, Salt Lake City, UT) in the hand area of motor cortex (M1) of each monkey (**Fig 1c**; monkey G: right, monkey P: left). Neural signals were collected using a Cerebus system (Blackrock Microsystems, Salt Lake City, UT). For the in-lab recordings, the neural signals were amplified by a Cereplex-E headstage. For the in-cage recordings, the neural signals were amplified, digitized at 30 kHz, and transmitted by a Cereplex-W wireless headstage. The neural signals were then digitally band-pass filtered (250 – 5000 Hz). Spikes were detected using a threshold set at -5.5 times the root-mean-square (RMS) amplitude of the signal on each channel, and the time stamp and a 1.6 ms snippet of each signal surrounding the threshold crossing were recorded. We used multiunit threshold crossings on each channel instead of well isolated single neurons in all our data analyses. We applied a Gaussian kernel (S.D.: 100 ms) to the spike counts in 50 ms, non-overlapping bins to obtain a smooth firing rate as function of time for each channel. We excluded channels with an average firing rate < 0.5 Hz during either in-lab or in-cage movements.

We also implanted intramuscular electromyographic (EMG) leads in 23 arm, forearm, and hand muscles in the left arm of monkey G and in the right arm of monkey P, contralateral to the M1 implant. For the in-lab recording sessions of monkey G, we collected EMG signals using a multi-channel differential amplifier and the analog input channels of the Cerebus system. For the in-cage sessions of monkey G and all sessions of monkey P, we collected EMG signals using a micro multi-channel amplifier (RHD2132, Intan Tech., Los Angeles, CA) and a wireless transmitter (RCB-W24A, DSP Wireless Inc., Haverhill, MA). The micro amplifier and wireless transmitter were both placed in a backpack on the monkey’s jacket. The EMG signals were amplified, band-pass filtered (4-pole, 50 - 500 Hz), and sampled at 2000 Hz. The EMGs were subsequently digitally rectified and low-pass filtered (4-pole, 10 Hz, Butterworth), and subsampled to 20 Hz (50 ms bins) to match the neural spike counts.

For each monkey, we divided the neural and EMG data collected for a given task into smaller datasets of non-overlapping data segments of 75 seconds duration. We chose 75 seconds based on simulations^24^ and on a preliminary analysis (**Supplementary Fig 1**) that assessed the effect of the amount of temporal data on dimensionality estimates. All neural and EMG data channels were normalized to their 95^th^ percentile value.

### Embedding and intrinsic dimensionality estimation algorithms

We used two algorithms for dimensionality estimation: Parallel Analysis^31–33^ and Two Nearest-Neighbors^34^. These methods were selected due to their superior performance in evaluating the dimensionality of simulated linear and nonlinear manifolds, respectively^24^.

Parallel Analysis (PA) is an embedding dimensionality estimator based on the eigenvalues of the covariance matrix for the dataset^31^. Unlike other embedding dimensionality estimators that rely on eigenvalue computation, PA does not rely on a predetermined variance threshold, but rather relies on counting the number of eigenvalues that exceed their respective values in a null distribution. This method has been shown to be an accurate estimator of embedding dimensionality in many fields^35^.

To estimate the embedding dimensionality of a given *M*-by-*N* dataset with *M* samples of *N* recorded signals, the PA algorithm proceeds as follows. First, it repeats the following process *K* times: for each feature, the data are independently shuffled using a random permutation along the corresponding column, the temporal axis; this shuffling breaks the correlation across features. Then, the eigenvalues of the features covariance matrix for the shuffled data are obtained and sorted from largest to smallest. A null distribution is created for each eigenvalue from the *K* shuffles. We used *K* = 200 repetitions and fixed the random number generator seed for reproducibility.

Next, we computed the 95^th^ percentile from the null distribution of each eigenvalue; this serves as a significance threshold. Finally, we counted the number of original eigenvalues that were larger than their respective 95^th^ percentile null distributions. This number was *D*_*PA*_, the embedding dimensionality estimate obtained by PA. While the 95^th^ percentile cutoff was an arbitrary choice, PA is quite robust to a reasonable choice of threshold to characterize significant deviations to the null, noise-based eigenvalue distribution^35^. In contrast, the usual practice of choosing a threshold for the variance-accounted-for (VAF) in a PCA-based approach often results in significant changes in the estimate of the embedding dimensionality when the threshold is changed by a few percentage points, such as choosing a 90^th^ instead of a 95^th^ cutoff.

To estimate the intrinsic dimensionality, we used Two Nearest-Neighbors (TNN), an estimator based on the local adjacency of data points^34,36,37^. Specifically, this method is based on the ratio *μ* of distances to the second vs the first nearest neighbors of a given point. The second-to-first nearest neighbors distance ratio is Pareto distributed with a unitary scale parameter and a shape parameter equal to the intrinsic dimensionality^38^. The intrinsic dimensionality can be approximated using the following equation:

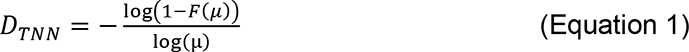

Here, F(*μ*) is the empirical cumulative distribution of the ratio of second-to-first nearest neighbor distances for each data point. In this study, we used the publicly available python package called `scikit-dimension`^39^ to compute the TNN estimate of intrinsic dimensionality.

### Denoising algorithms

Noise is a confounding factor for both intrinsic and embedding dimensionality estimates^24,29,30^. Random noise across channels will lead to increased dimensionality estimates that might even approach the total number of recorded signals. To address this issue, we implemented two denoising approaches that rely on an initial estimate of an upper bound dimensionality *D* = *D*_*PA*_ obtained using Parallel Analysis (PA).

The first approach, PCA denoising, is a linear method based on Principal Component Analysis (PCA, **Fig 2a**). After determining the value of *D* using PA, we used the *D* leading principal components to reconstruct the original data. This approach assumes that most of the noise is present in the low-variance principal components that have been discarded.

**Figure 2:**
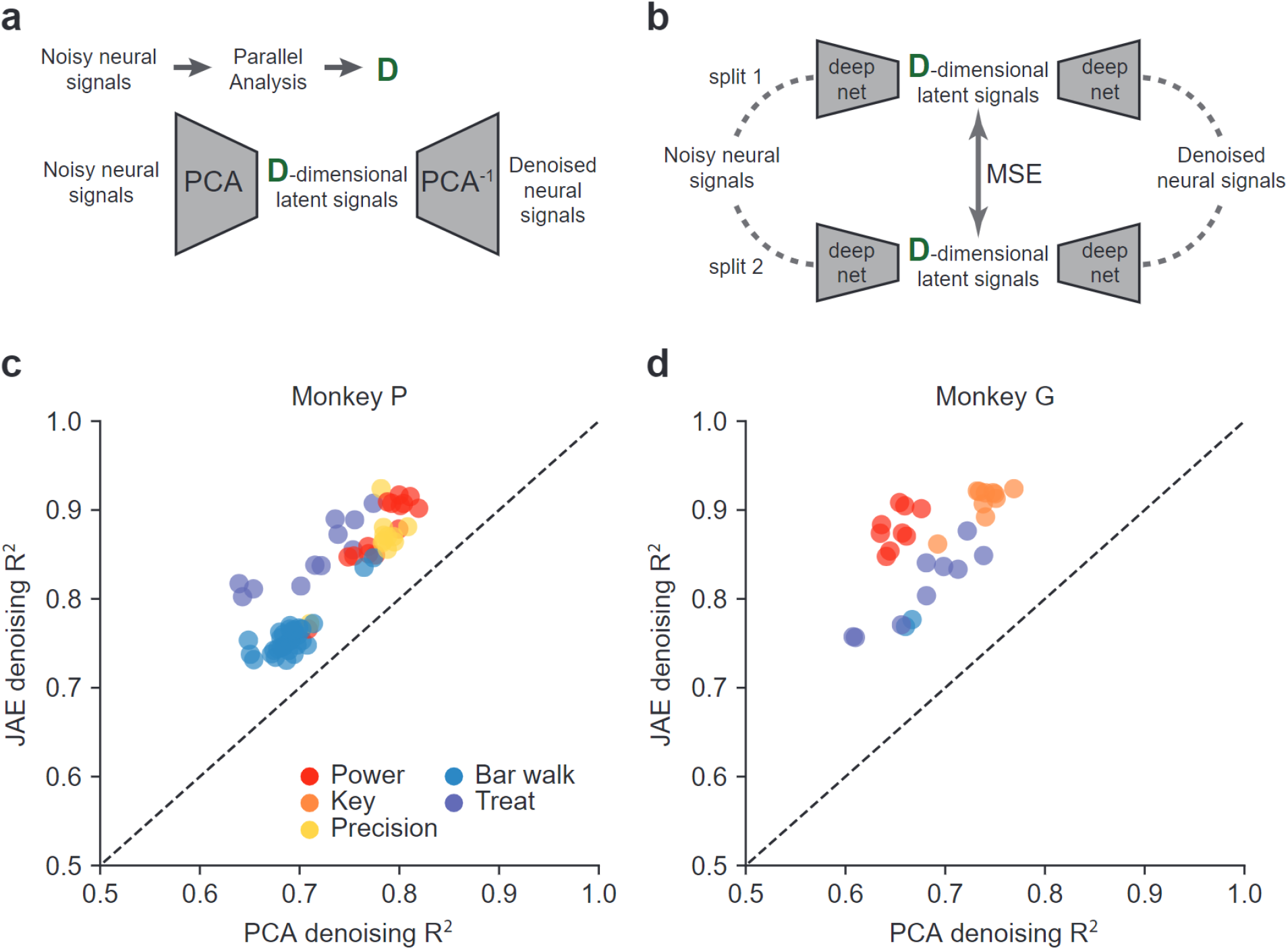
Denoising the neural signals. Two methods were employed to denoise neural signals: a PCA based approach and a deep neural network-based approach called JAE. a) In the PCA based denoising approach, noisy neural signals were projected to the leading *D* principal components, with *D* determined by Parallel Analysis. The denoised neural signals were the reconstructions from the *D*-dimensional bottleneck. b) For the deep neural network-based denoising approach we used the joint autoencoder (JAE). Noisy neural signals were randomly divided into two subsets. Each subset was nonlinearly projected to a *D*-dimensional latent space using two parallel deep networks. *D* was again determined using Parallel Analysis. The mean squared error (MSE) loss used to train the network forced the *D*-dimensional latent representations across the parallel networks to match. Denoised neural signals were obtained from the reconstructions based on the activity in the *D*-dimensional bottlenecks. c) Comparison of reconstruction accuracies between PCA and JAE denoising approaches for Monkey P. Each circle represents the R^2^ performance of each denoising method for a single dataset. Warm colors indicate the constrained laboratory tasks. Cold colors indicate the unconstrained cage tasks. d) Same as in panel c but for Monkey G.

The second approach, Joint Autoencoder (JAE) denoising, is a neural network-based method (**Fig 2b**). We divided the noisy neural signals with *N* features into two disjoint subsets, each including *N*⁄2 features, and used the compressive halves of two autoencoders to map each subset into a *D*-dimensional subspace. The reconstructed versions of the subsets of dimension *N*⁄2 resulted from the expansive halves of the respective autoencoders. The JAE was trained using a cost function that minimized the mean-squared error associated with the reconstructions of each of the two subsets, as well as the mean-squared error between the two intermediate *D*-dimensional latent signals. This approach assumes that each subset contains the necessary information to identify the underlying *D*-dimensional signals while the noise components are common across the two subsets.

To evaluate the performance of the denoising algorithms, we calculated the coefficient of determination (*R*^2^) between the denoised and noisy neural signals (**Fig 2c and 2d**). We validated these denoising approaches in a previous study based on simulated neural datasets with known levels of noise^24^.

### Robustness of the dimensionality estimates relative to the ambient dimensionality of the underlying neural space

Accurately interpreting the dimensionality of neural data requires consideration of the amount of data used for dimensionality estimation. Prior studies have shown that the amount of data used, measured by both the number of samples *M* and number of features *N*, can affect the accuracy and precision of the dimensionality estimates^1,23,24,30,40^. To ensure stable dimensionality estimates, we determined that datasets roughly one minute in length were sufficient for our analyses (see **Supplementary Figure 1**).

Similarly, we conducted analyses to determine the robustness of our estimates with respect to the number of neurons. Specifically, we aimed to assess whether our dimensionality estimates reached an asymptote as the number of neurons increased, indicating reliable estimates. We gradually sampled increasing numbers of neurons and repeatedly computed the dimensionality estimates 20 times for each number. We sampled 5, 10, 15, 20, 25, 30, 35, and the maximum number of recorded neurons for each dataset. If the estimates reached an asymptote, we concluded that additional neurons would not contribute meaningfully to the estimated dimensionality, indicating that our dimensionality estimates were robust. Conversely, if the estimates continued to increase with the number of neurons, this suggested that more neurons would be necessary to obtain reliable estimates.

### Quantifying the extent of nonlinearity

We used two measures to compute the extent of nonlinearity in the neural recordings; these measures arise from two complementary perspectives. The first measure, that we called *manifold nonlinearity index*, focused on the nonlinearity of the geometry of the neural manifold. The second measure, that we called *local flatness index*, focused on how much of the population activity is concentrated in approximately linear regions within the manifold.

To compute the manifold nonlinearity index, we used the ratio of embedding-to-intrinsic dimensionality of the neural manifold^24^. The manifold nonlinearity index is similar to the dimensionality gain metric used in an artificial neural network study that sought to extract predictive latent signals^41^.

To compute the local flatness index, we compared the Euclidean and geodesic distances between every pair of data points in the state space of each dataset (**Supplementary Figure 2**). Euclidean distance is the length of the shortest straight line between two points, while geodesic distance takes into account the curvature of the neural manifold by measuring the length of the shortest path that lies within the manifold and connects the two points. To calculate the geodesic distance from one point to all others, we started from a nearest-neighbor cloud limited to 200 samples. Distances to those 200 closest activity patterns were computed along the straight lines joining the central point to its neighbors within the cloud. It is from these short linear segments that global geodesics are constructed.

A large discrepancy between the Euclidean and geodesic distances indicates that the population activity explores curved, nonlinear regions within the manifold. A higher degree of overlap suggests that the neural activity is largely confined to nearly linear regions within the manifold. To determine the extent to which the neural activity evolves in a linearizable region of the neural manifold, we computed the empirical distribution of both Euclidean and geodesic distances for each dataset. We used 20 bins to cover the full range shared between these two distance distributions; we then converted these distributions to normalized densities. Within each bin, the smaller of the two density values, multiplied by the bin size and summed over all bins provides the value for the local flatness index. **Supplementary Figure 2** shows a graphical illustration of how we computed the local flatness index for a randomly chosen bar walk dataset from Monkey P.

### Computing the activity on the low-dimensional neural manifolds

To project the population neural activity onto the low-dimensional neural manifolds, we employed two different techniques, each with different underlying assumptions about the linearity of the data: Principal Component Analysis (PCA) and a nonlinear autoencoder. PCA is a widely used linear technique based on finding the orthogonal directions in neural space that correspond to maximum variance in the data. In contrast, the nonlinear autoencoder is a neural network-based technique that can capture complex nonlinear relationships in the data and is especially useful for projecting population activity onto the low-dimensional nonlinear neural manifolds. The autoencoder consisted of five feedforward layers (input, first hidden, bottleneck, second hidden, and output layers) with ReLU activation and was trained using the ADAM optimizer on the reconstruction error, the mean-squared error between output and input; default training parameters were used, and training lasted for 50 epochs.

For both techniques, we used the TNN estimates of the intrinsic dimensionality to determine the manifold dimensions. For PCA, this meant that we retained a number of leading principal components equal to *D*_*TNN*_ rounded to the nearest integer. For the autoencoder, we made the number of neurons in the bottleneck layer equal to *D*_*TNN*_ rounded to the nearest integer.

### Decoding electromyograms (EMGs)

Our setup allowed for the simultaneous recording of neural signals from the primary cortex and EMG signals from muscles as the monkeys engaged in various motor behaviors. We used linear regression to decode EMG signals from neural signals; this basic, interpretable, and linear method is widely used in the field of brain-computer interfaces^42,43^. Each set of EMG signals was decoded using three types of inputs: all available neurons, latent signals obtained from PCA, and latent signals obtained from the bottleneck layer of the autoencoder. We used five-fold cross validation for each approach and reported all five test folds for a given neural-to-EMG dataset pair.

### Statistical analyses

We reported our computations in the mean ± standard deviation format. We used Welch’s t-test for statistical comparisons when the distribution of the statistic of interest was normal. We used Wilcoxon rank sum test in the case when the normality assumption was violated (**Supplementary Fig 3**). When we compared EMG decoding accuracies from PCA embeddings, AE embeddings, and all available neurons, we used repeated measures ANOVA (**Fig 6**). We reported the *p*-values obtained from the statistical test in the figure legends and used Bonferroni correction for multiple comparisons where applicable.

### Ethics statement

All surgical and experimental procedures that yielded the datasets comprising of multi-electrode neural recordings and intramuscular electromyogram (EMG) signals from non-human primates were approved by the Institutional Animal Care and Use Committee of Northwestern University. The two monkeys were monitored daily to ensure their well-being and health. Their diet consisted of a standard laboratory animal diet supplemented with fresh fruits and vegetables to provide optimal nutrition. Additionally, the monkeys were provided with access to various types of enrichment to promote mental stimulation and overall well-being.

## RESULTS

### Reducing the effects of noise using linear and nonlinear methods

Both neurons and neural recordings are noisy^44^, which can cause an overestimation of the dimensionality^23,24,34,45^ of recorded neural activity. To mitigate the effect, we first denoised the recorded neural signals using two approaches that differed by their assumption about the linearity of the manifold to which the data is mostly confined. The linear denoising method was based on Principal Component Analysis (PCA, **Fig 2a**). The nonlinear denoising method was a neural network-based approach called the Joint Autoencoder (JAE, **Fig 2b**).

For every data set for both monkeys, denoising the data using JAE yielded a higher reconstruction accuracy (R^2^) than denoising the data using PCA (**Fig 2c and 2d**). For monkey P and for the power and precision grasp tasks, PCA yielded a reconstruction accuracy of 0.78 ± 0.03 (mean ± standard deviation) and 0.78 ± 0.02, whereas JAE yielded 0.88 ± 0.04 and 0.87 ± 0.04, respectively. The reconstruction accuracies for the bar walk and treat grasp tasks were 0.69 ± 0.02 and 0.71 ± 0.05 with PCA, and 0.76 ± 0.02 and 0.85 ± 0.04 with JAE. We observed a similar trend for Monkey G. For the power and key grasp tasks, reconstruction accuracy using PCA was 0.65 ± 0.01 and 0.74 ± 0.02, whereas the reconstruction accuracy using JAE was higher at 0.88 ± 0.02 and 0.91 ± 0.02. For the unconstrained tasks, reconstruction accuracies using PCA and JAE were 0.66 ± 0.01 and 0.68 ± 0.05, and 0.77 ± 0.01 and 0.81 ± 0.04, respectively. For the reminder of the analyses, we denoised the neural signals using the JAE approach due to its consistently superior performance over PCA.

### Dimensionality estimates of denoised neural signals

Our next goal was to compute the dimensionality of the denoised signals. We applied two dimensionality estimation algorithms: PA and TNN. We used PA to estimate the embedding dimensionality and TNN to estimate the intrinsic dimensionality of the neural signals. These methods were chosen based on their assumption about linearity and superior performance to alternatives^24^.

Both embedding and intrinsic dimensionality estimates were slightly higher for unconstrained behaviors compared to constrained behaviors for both Monkey P and Monkey G (**Fig 3**). Dimensionality estimates from PA were almost always higher than those from TNN. For Monkey P, the PA dimensionality across the constrained laboratory tasks increased from 7.28 ± 1.67 to 10.40 ± 2.54 across the unconstrained tasks (**Fig 3a**). Similarly for Monkey G, the PA dimensionality across constrained and unconstrained tasks was 5.84 ± 0.78 and 9.27 ± 1.62, respectively (**Fig 3b**). We observed the same trend in TNN dimensionality across the two task settings. TNN dimensionality estimates for Monkey P increased from 5.01 ± 0.39 to 6.5 ± 0.58 from the constrained to unconstrained tasks (**Fig 3c**). Similarly for Monkey G, TNN estimates went from 4.72 ± 0.18 to 5.70 ± 0.27 (**Fig 3d**). In summary, neural manifolds for all tasks were consistently much smaller than the total number of sampled neurons, which defines the dimensionality of the empirical neural space that contains these task-specific manifolds. We note that both the intrinsic and embedding dimensionalities were slightly higher for unconstrained tasks.

**Figure 3:**
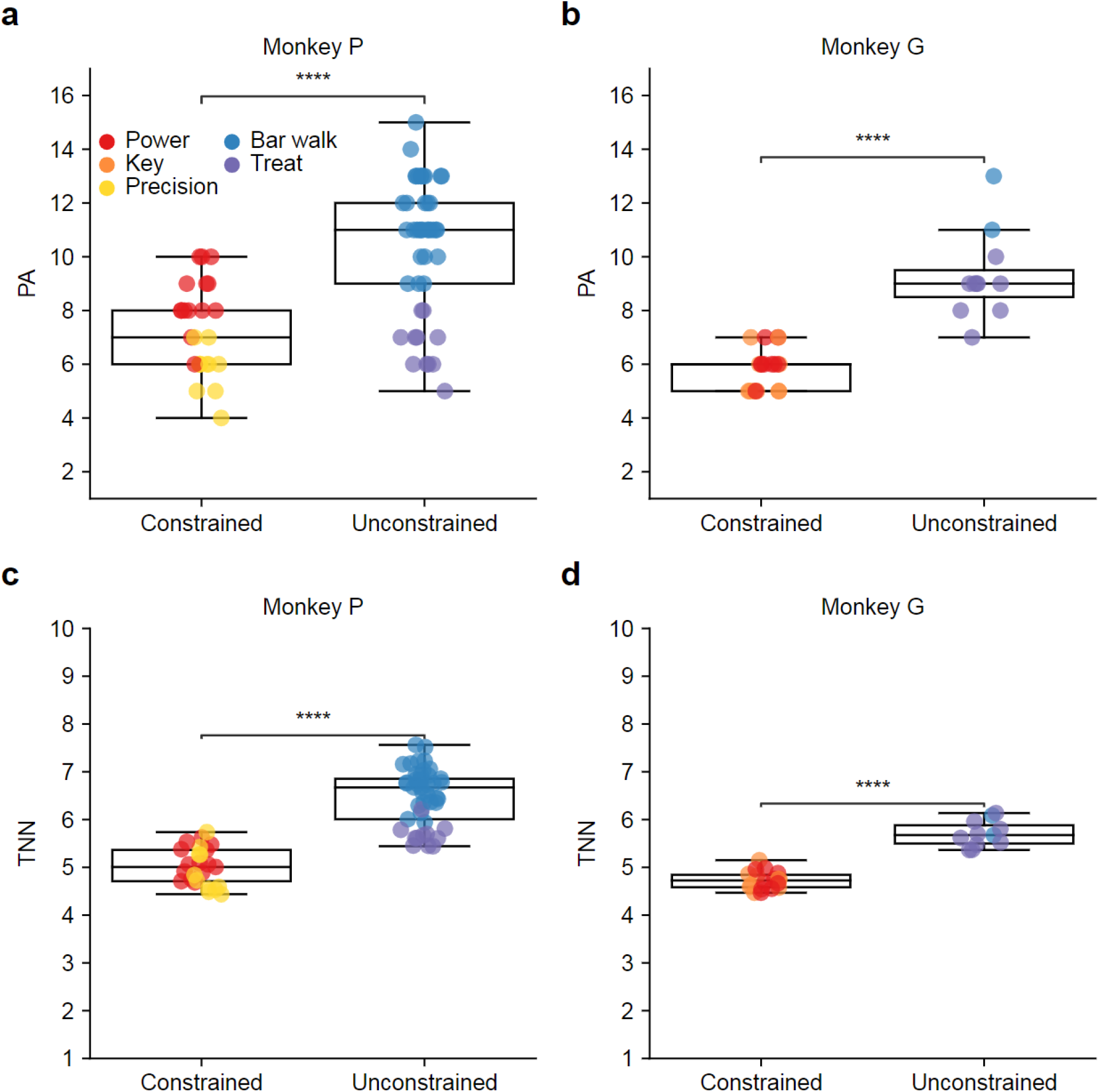
Estimating the embedding and intrinsic dimensionalities of neural manifolds. We applied Parallel Analysis (PA) and Two-Nearest Neighbors (TNN) to estimate the embedding and intrinsic dimensionality, respectively. a) Comparison of the PA estimates for constrained and unconstrained behaviors for Monkey P (p ≈ 0). Each circle represents the dimensionality estimate from a single dataset. Warm colors indicate constrained behaviors in the laboratory. Cold colors indicate unconstrained behaviors in the cage. b) Same as in panel a but for Monkey G (p ≈ 0). c) Comparison of the TNN estimates across constrained and unconstrained behaviors for Monkey P (p ≈ 0). d) Same as in panel c but for Monkey G (p ≈ 0).

We asked if the small but significant increase in the dimensionality of neural manifolds associated with unconstrained tasks is a reflection of increased task complexity^1,25^, and used the intrinsic dimensionality of EMG signals as a proxy for task complexity (**Supplementary Fig 3**). The increase in EMG dimensionality when comparing unconstrained to constrained tasks was also small but significant, and particularly noticeable in some instances of unconstrained tasks for monkey G (**Supplementary Fig 4**).

### Investigating the extent of nonlinearity in the neural manifolds

To quantify the degree of nonlinearity in the geometry of neural manifolds, we computed the *manifold nonlinearity index* (see Methods). On average, the manifold nonlinearity index was slightly higher for the unconstrained tasks than for the constrained tasks, but rarely exceeded two (**Fig 4a and 4b**). On average, the manifold nonlinearity index was 1.45 ± 0.29 (Monkey P) and 1.24 ± 0.16 (Monkey G) for the constrained tasks, and 1.58 ± 0.28 (Monkey P) and 1.63 ± 0.26 (Monkey G) for unconstrained tasks, respectively. The larger manifold nonlinearity index for Monkey P was not statistically significant.

**Figure 4.**
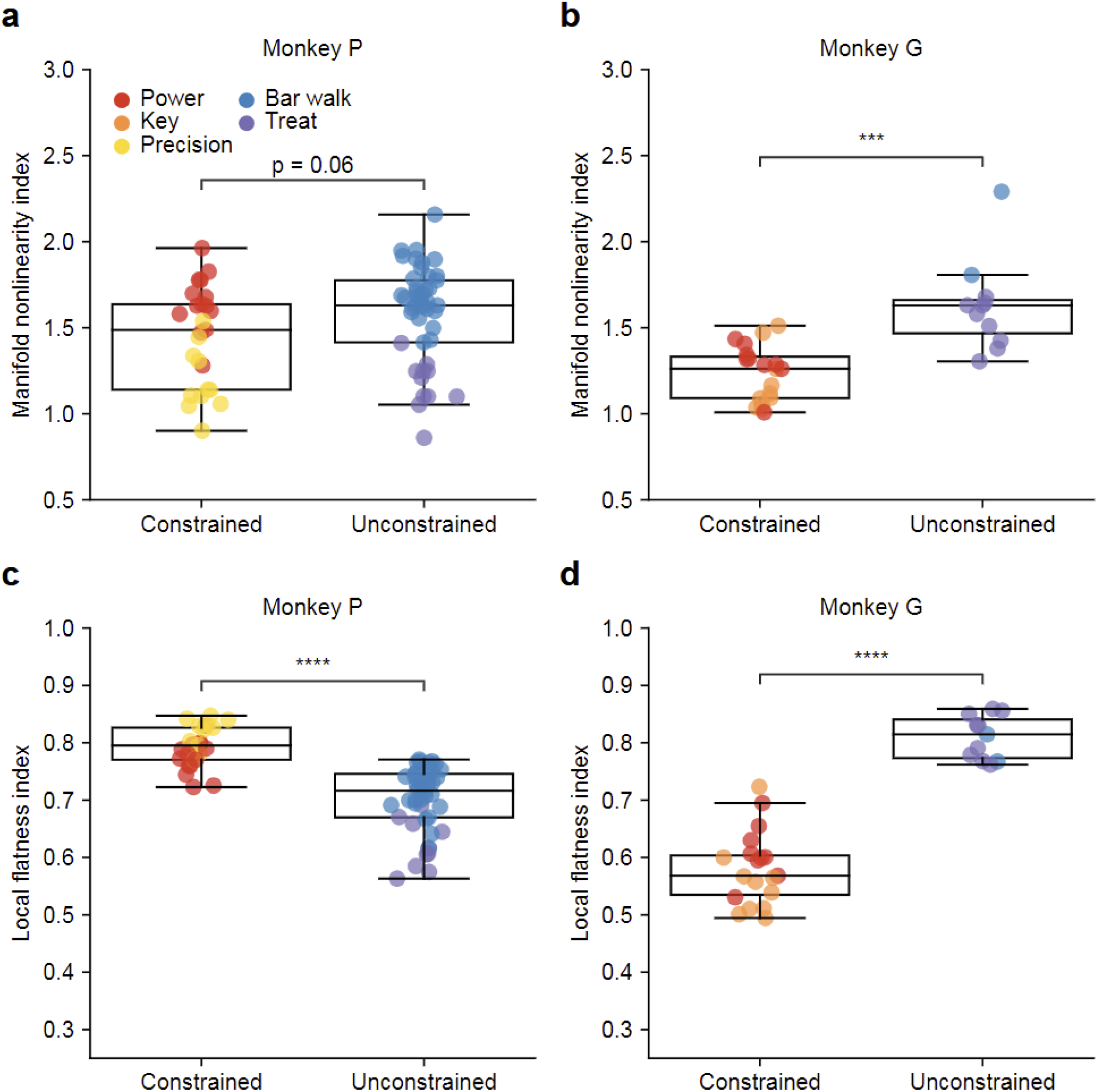
Manifold nonlinearity index and local flatness index of neural manifolds. a) Manifold nonlinearity index, the ratio of the embedding (PA) to intrinsic (TNN) dimensionality, for all datasets for Monkey P. Each circle represents the manifold nonlinearity index for a single dataset. Warm colors indicate constrained behaviors in the laboratory. Cold colors indicate unconstrained behaviors in the cage (p = 0.06). b) Same as in panel a but for Monkey G (p = 0.0005). c) Local flatness index, measured by the fractional overlap between the distributions of Euclidean and geodesic distances (see Methods, Supplementary Figure 2), for Monkey P (p ≈ 0). d) Same as in panel c but for Monkey G (p ≈ 0).

The manifold nonlinearity index only partially elucidates the nature of nonlinearity of the neural manifold, because it does not reveal how neural activity samples the manifold. To complement it, we computed the local flatness index, which quantifies the extent to which the population activity samples linear regions within the manifold (see Methods). The approach is based on comparing all pairwise Euclidean and geodesic distances for each dataset. (**Fig 4c and 4d**). The overlap between the distributions of geodesic and Euclidean distances highlights the degree of linearity in the explored regions of the manifold: an overlap greater than 0.50 indicates that most of the data lie in regions of the manifold that are well approximated by the tangent linear subspace (**Supplementary Fig 2**).

For Monkey P, the constrained tasks had a local flatness index of 0.79 ± 0.04 and unconstrained tasks had a local flatness index of 0.70 ± 0.06. For Monkey G, local flatness was 0.58 ± 0.06 for constrained and 0.81 ± 0.04 for the unconstrained tasks (**Fig 4c and 4d**). The differences for both monkeys were statistically significant, despite being in opposite directions. We notice the low value of the local flatness index for Monkey G when executing constrained tasks, and hypothesize that this effect is likely associated with the adoption of distinct task execution strategies by Monkey G in the laboratory, as illustrated in **Supplementary Fig 3b**. For this hypothesis to be confirmed, we would need to analyze and compare the fine details of task execution, an analysis beyond the scope of this work. To summarize these findings the aggregate results in **Fig 4c and 4d** show that the local flatness index never fell below 0.5 in any constrained or unconstrained scenario; this indicates a substantial overlap between the distributions of geodesic and Euclidean distances for both monkeys. Results for the local flatness index establish that for the tasks investigated here the neural population dynamics visited states mostly confined to nearly linear regions within slightly nonlinear neural manifolds.

### Number of neurons required for stable dimensionality estimates

One critical consideration for interpreting dimensionality is the amount of data needed for dimensionality estimates; previous studies show that the amount of data can affect the accuracy of the estimates^24,40^. We determined that datasets roughly one minute-long were sufficient for stable dimensionality estimates **(Supplementary Figure 1).**

Another factor contributing to the reliability of dimensionality estimates is the number of recorded neurons. How robust are the dimensionality estimates with respect to the number of neurons used in analysis? To answer this question, we assessed whether the dimensionality estimates reached an asymptote as the number of neurons used for analysis increased. An asymptotic saturation of the estimated dimensionality would signal reliable dimensionality estimates.

We found that for both monkeys the nonlinear TNN method saturated as the number of neurons increased (**Fig 5a and 5c**). The TNN dimensionality estimates saturated at roughly 20 to 35 sampled neurons for all tasks and monkeys. In contrast, the linear method PA yielded dimensionality estimates that continued to increase as the number of neurons used to estimate manifold dimensionality increased (**Fig 5b and 5d**).

**Figure 5:**
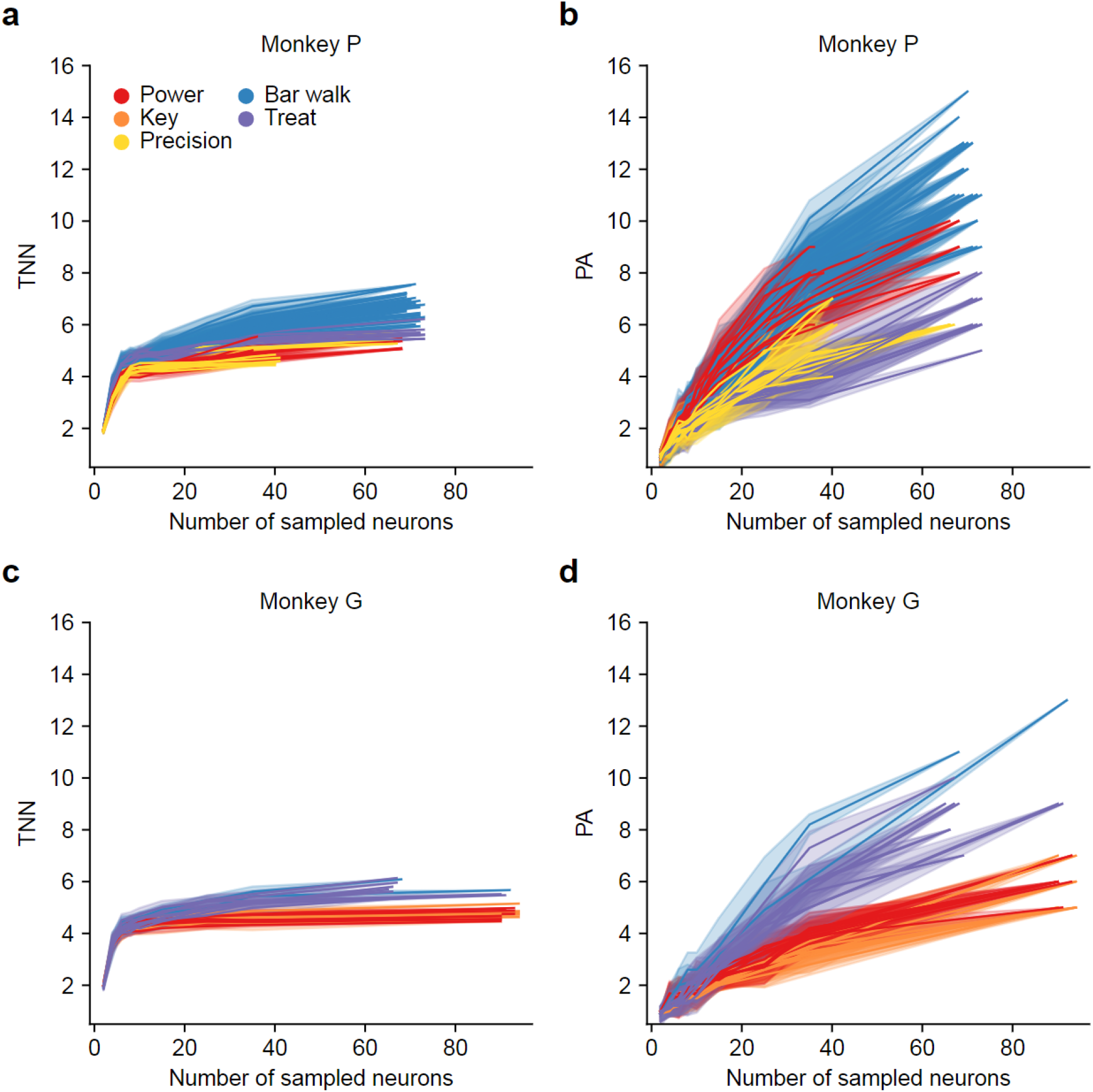
Number of neurons required for stable dimensionality estimates. We progressively increased the number of neurons included in the dimensionality estimation. We randomly subsampled, with ten repetitions, a given number of neurons from the total number of neurons available in each dataset. Solid lines indicate the average dimensionality estimate. Shaded regions indicate the standard deviation. a) TNN estimates from for Monkey P. b) PA estimates for Monkey P. c) Same as in panel a, but for Monkey G. d) Same as in panel b, but for Monkey G.

### Decoding EMGs from low-dimensional latent activity

Finally, we compared how well the neural manifolds represented behavior by decoding EMGs from the activity within the low-dimensional neural manifolds obtained with both PCA and a nonlinear autoencoder (**Fig 6**). We compared the accuracy of these decoders to that of decoders based on the activity of all recorded neurons. In both scenarios, we reported the *D*_*TNN*_ estimates of the intrinsic dimensionality of the EMG signals and used this dimensionality as the number of leading latent variables to be used as inputs to a neural-to-EMG decoder. This approach allowed us to directly quantify the extent to which muscle-related information lives in a linear hyperplane that approximates a slightly nonlinear neural manifold of *D*_*TNN*_ dimensions. Examples of actual and decoded EMG signals based on all recorded neurons, linear latent variables, and nonlinear latent variables are shown in **Supplementary Fig 5** for five different muscles. We also showed the EMG decoding performance from progressively increasing neural manifold dimensionality in **Supplementary Fig 6**.

**Figure 6:**
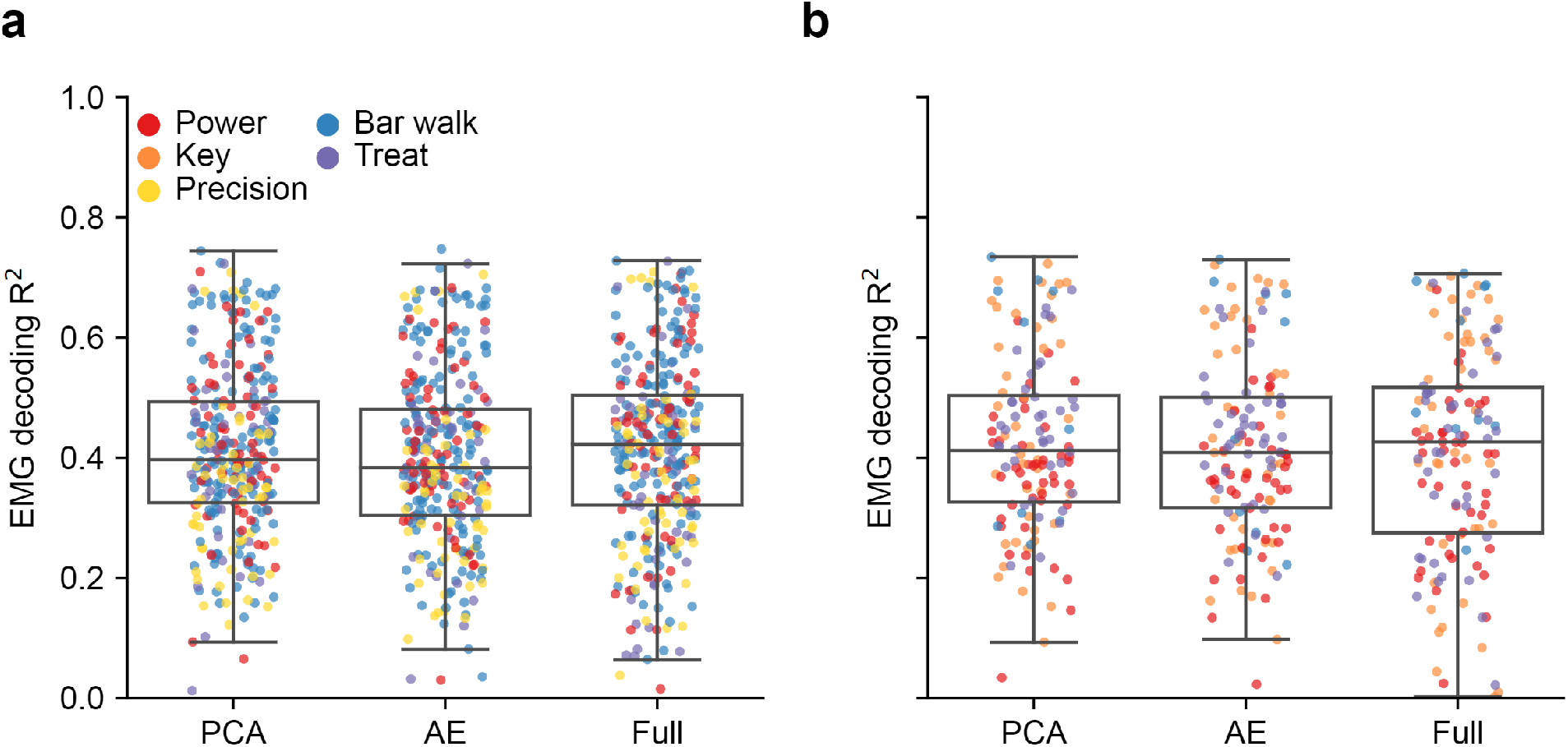
Decoding EMGs from low dimensional embeddings and all available neurons. We used linear regression to decode EMGs from denoised neural signals (see Methods for denoising). We either used PCA embeddings (PCA), AE embeddings (AE), or the full-dimensional neural signals (Full) as inputs to linear neural-to-EMG decoders. PCA and AE provided linear and nonlinear latent variables, respectively. The dimensionality of the latent representations followed from the TNN estimate for each dataset. Each dot represents the decoding accuracy from one of the five test folds of decoding. Different colors represent different tasks. Repeated measures ANOVA was not significant between the three conditions for two monkeys (p = 0.59). a) EMG decoding accuracies for Monkey P. b) Same as in panel a, but for Monkey G.

In all datasets, the accuracies obtained when decoding EMGs from all recorded neural signals and from the low-dimensional latent variables were not significantly different. Importantly, linear decoding from the activity of PCA and autoencoder latent variables restricted to *D*_*TNN*_ dimensions yielded similar accuracy–ANOVA with repeated measures was not significant, p = 0.59. For Monkey P, the average test-fold EMG decoding accuracy using PCA, autoencoder, or all recorded neural signals was (mean ± standard deviation) 0.41 ± 0.14, 0.40 ± 0.15, and 0.41 ± 0.15 (**Fig 6a**). The corresponding results for Monkey G were 0.42 ± 0.15, 0.42 ± 0.15 and 0.41 ± 0.17(**Fig 6b**). These results indicate that there was no difference in EMG decoding from the activity of all recorded neurons or from the latent variables that characterize the low-dimensional neural manifolds. Additionally, the similar EMG decoding accuracy from *D*_*TNN*_ latent variables obtained from PCA and nonlinear autoencoder embeddings indicates that the information encoded in the motor cortex relevant to activating muscles exists in a mostly linear subspace of dimension *D*_*TNN*_ (**Supplementary Fig 5 and 6**).

## DISCUSSION

Neural dimensionality has emerged as an important concept for understanding the underlying dynamics and computation abilities of populations of neurons. The dimensionality of the primary motor cortex (M1) manifolds associated with specific tasks has been found to be low in numerous studies limited to constrained laboratory behaviors^2,9,24,27,46^. In this work, we computed the intrinsic and embedding dimensionalities of M1 recordings as monkeys engaged in unconstrained tasks in their home cage. Our primary finding was that both the intrinsic and embedding dimensionality of M1 signals were slightly higher than for tasks performed with a single arm, while seated in a primate chair. Although the manifolds associated with unconstrained tasks were found to have slightly higher dimensionalities, these dimensionalities were still very low. In addition, although we found signatures of nonlinearity in the M1 neural manifolds, most of the activity was confined to nearly linear evolved regions within the neural manifolds. Finally, the accuracy of linearly decoded EMGs from the low-dimensional latent variables that characterize the manifolds matched closely the accuracy of EMGs decoded from all recorded neurons, both in cage and in lab.

### Linking neural dimensionality to neural computation and processing

There are different definitions and interpretations of dimensionality. We adopted those recently put forward by Jazayeri and Ostojic, who distinguish between intrinsic and embedding dimensionalities^23^. While the intrinsic dimensionality quantifies the number of independent latent variables encoded by a neural population, the embedding dimensionality plays a role in understanding how the latent variables are processed and transmitted. Together, the analysis of both intrinsic and embedding dimensionalities illuminates how neural systems encode and process information.

Intrinsic dimensionality refers to the actual dimensionality of a nonlinear manifold to which the population neural activity is mostly confined; it may include different classes of information, such as sensory inputs, motor outputs, and other latent variables that correspond to learned experiences and expectations^23^. Since different brain regions selectively represent these classes, the intrinsic dimensionality of a representation in a particular brain area may be strongly associated with one class and weakly with another. For example, the intrinsic dimensionality of latent signals in early sensory areas like the olfactory cortex and primary visual cortex is associated with the representation of incoming stimuli such as chemical odors^47^ or visual gratings^48^, respectively. On the other hand, latent signals from the primary motor cortex are more closely associated with motor output^2,6,49,50^ and less with future expectations^51^. The intrinsic dimensionality of manifolds in the primary motor cortex would therefore be expected to be closely associated with motor output variables such as kinematics or muscle activation.

Embedding dimensionality refers to the dimensionality of the minimal Euclidean space sufficient to fully contain the nonlinear manifold. Although the recurrent dynamics within a given brain area might lead to nonlinear manifolds best characterized by their intrinsic dimension, it is their embedding dimension that best characterizes the signals to be communicated through linear readouts. This observation has led to the view that the embedding dimension reflects how the latent variables are processed for communication to other neural areas^23^. However, the computational principles associated with the embedding dimensionality are not yet fully understood. What determines whether the embedding dimensionality is high or low? We have argued that a higher degree of manifold nonlinearity will require a larger embedding dimension. But how does this concept connect to the question of communication across areas? One hypothesis is that the neural code is linearly relayed to other brain areas or to the periphery^52–56^.

In this view, different linear decoders can selectively relay different task relevant information to different downstream areas without interference, a property called mixed selectivity^57,58^. A brain area that relays distinct sets of information to several downstream areas would require the latent variables to be embedded into a higher-dimensional Euclidean subspace to facilitate mixed selectivity. Mixed selectivity thus necessitates a high embedding dimensionality^53^. As an example of this correspondence, prefrontal and primary visual cortical neurons have been reported to have a very high embedding dimensionality^52,57,59^; these observations fit well with the mixed selectivity hypothesis. In contrast, a brain area that does not require extensive mixed selectivity, such as the primary motor cortex, would generate latent variables that only require a relatively low embedding dimensionality, an observation consistent with many reports on M1 manifolds^2,9,24,27,28,60,61^.

### Low dimensionality of M1 transcends task constraints

Thus far, the low dimensionality of M1 manifolds has only been investigated in the context of constrained laboratory settings. It is unclear whether the low dimensionalities observed in M1 truly reflect some intrinsic computational property of M1 or whether they are a byproduct of the constraints in stereotyped, repeated laboratory tasks. To begin to elucidate this question, our approach was to obtain and analyze a rich collection of datasets corresponding to two distinct settings: laboratory and cage. These two settings captured different levels of constraint. In the laboratory setting, we placed the monkeys in a primate chair with restraints such that they were only able to move the hand contralateral to their neural implant in specific, well-instructed trial segments. In the cage setting, the monkeys had more freedom to move around the perch bars and could grab small treats from the experimenters as they pleased. We did not provide any instructions on how or when to perform the tasks in the cage; the monkeys could take as much time as they needed and execute these tasks in a non-stereotyped manner. Our results show that the low dimensionality of the primary motor cortex is largely independent of task setting and constraints. Both the intrinsic and embedding dimensionalities of M1 were low in both constrained laboratory and unconstrained cage settings, with only slightly higher dimensionality estimates for the unconstrained tasks (**Fig 3**). Thus, we hypothesize that the low dimensionality of M1 may not be a byproduct of constrained movements, but rather reflects its computational strategy. Our results, obtained through the lens of dimensionality, fit in with recent evidence of context independence in M1^15^. In contrast, recent evidence from mouse cerebellar parallel fiber recordings showed a roughly four-fold increase in embedding dimensionality from constrained limb-actuated lever tasks to spontaneous standing, running, or whisking tasks^62^. The difference between M1 and cerebellar results, though from a different species, highlights that different brain areas differentially process task constraints and context.

### M1 representations are nonlinear, but only slightly

Task-specific M1 representations were slightly nonlinear for all tasks and in both settings. For a given bottleneck dimension *D*, the nonlinear JAE denoising algorithm consistently resulted in a better reconstruction of neural signals than PCA (**Fig 1**). The manifold nonlinearity index was between one and two for all datasets (**Fig 4**). Finally, the local flatness index was, on average, around 0.7 when aggregated across all datasets. Although these results describe neural manifolds that were monkey and task specific, a general trend emerges: despite the slight degree of nonlinearity in the geometry of the neural manifolds, task-specific neural dynamics sampled mostly linear regions within the respective nonlinear manifolds (**Fig 4**). In other words, there was evidence of only mild nonlinearity in M1 manifolds associated with the motor behaviors analyzed here; in addition, the population activity was mostly confined to linear regions within these manifolds.

### Interpreting the low dimensionality and mild nonlinearity of M1

How should we interpret the low dimensionality and mild nonlinearity of M1 manifolds? One explanation is related to the computational tradeoff between generalizability and expressivity that has been studied in both artificial and biological neural networks^63–66^. Let’s first consider an extreme case: if the task-specific information encoded in a brain area must be selectively communicated to several different brain areas, latent signals ought to be confined to distinct, linearly independent subspaces, and would therefore require a high embedding dimension. Such an organization would facilitate linear readouts without interference and would therefore be more expressive^23,52,53,57^. The advantages of high-dimensional embeddings for linear readouts are well understood in artificial networks and widely used in the context of kernel methods and support vector machines^54,67,68^.

While there is evidence of some degree of expressivity in M1^3,6,8^, our findings of low-dimensional manifolds implies that mixed selectivity is lower in M1 than in some higher-order brain areas. For example, a brain area that must selectively relay the abstract variables relevant to decision making and ultimately leading to motor output, such as the dorsolateral prefrontal cortex, requires high mixed selectivity and has been reported to have a large embedding dimensionality^52,57^. The low dimensionality and mild nonlinearity of M1 indicates that such complex representations are unnecessary for M1. Instead, M1 exhibits a more generalized, low-dimensional representation that encodes different inputs into a small set of common activity patterns^52,53^. Thus, from a functional perspective, the low-dimensional and generalizable computational strategy of M1 facilitates the reliable generation of muscle commands that are largely unaffected by task constraints.

Another interpretation of the low dimensionality and mild nonlinearity of M1 neural population activity refers to the strength of recurrent connectivity in M1. The dynamical systems perspective highlights the existence of recurrent connections, but we do not know much about the strength of these connections^22^. Recent work on artificial and biological networks related the strength of recurrent connections to the embedding dimensionality of neural representations^69–73^. In this view, networks with low-rank connectivity matrices generate low-dimensional dynamics. While the body of work that relates recurrent connectivity structure to dynamics is in its infancy and warrants future studies, the low dimensionality of M1 is consistent with weak recurrent connections^74^.

### Decoding EMGs from low-dimensional manifolds

Recent theoretical work directly relates the dimensionality of M1 manifolds to the accuracy of movement parameter decoding^1,25^. In this view, the low dimensionality of M1 permits sampling far fewer than the millions of active neurons for accurate decoding. While this theoretical work was verified in laboratory settings, the relationship between low dimensionality and decoding accuracy had not been shown in more natural settings. We showed that decoding EMGs from latent variables works reasonably well in either context (**Fig 6, Supplementary Figures 5 and 6**), indicating that the proposed theoretical ideas also apply in unconstrained settings. Our results indicate that current hardware recording technology that samples only hundreds to thousands of neurons allows for the identification and description of neural manifolds associated with unconstrained tasks. Generalizable representations in M1 are low dimensional and in turn decodable even when our recording devices vastly undersample the active neurons in the motor cortex. These findings contrast with the recent findings about dimensionality and decoding accuracy in the prefrontal cortex (PFC). The high embedding dimensionality of PFC representations make it difficult to decode behavioral parameters form neural data acquired with current recording technology, despite moment-by-moment changes in neural activity during behavior^75^; in some cases, decoding behavior from PFC was barely above chance levels.

### Limitations

When the cage experiments were designed, we anticipated the unconstrained tasks to be more complex, and hoped to quantify this expectation through a higher intrinsic dimensionality of EMG signals. However, we failed to see the anticipated effect. For Monkey P, the EMG reconstruction accuracies with progressively increasing latent dimensionality of linear and nonlinear EMG manifolds were slightly lower for the treat grasp task than for the other behaviors, signaling higher task complexity (**Supplementary Figure 4a**). The reconstruction curves for bar walk overlapped with those of the constrained tasks. For Monkey G, the EMG reconstruction curves associated with both bar walk and treat grasp tasks appeared to be below those for constrained tasks (**Supplementary Figure 4b**). However, for all tasks and both monkeys, roughly 4 to 5 latent dimensions were sufficient to explain over 0.70 of the variance in EMG signals. Importantly, the TNN dimensionality of EMGs corresponding to constrained and unconstrained settings were similar, and between 2 and 6 for both monkeys (**Supplementary Figure 3**). These results indicate there was no significant difference in task complexity between constrained and unconstrained tasks, in contrast to our original expectations. Therefore, one limitation of our study was that all tasks that we tested were relatively simple, even the unconstrained tasks in the more naturalistic cage setting. A definitive comparison between neural manifold associated with simple versus complex tasks is still needed.

Interpreting the dimensionality and complexity of behaviors is difficult^76^. Earlier studies showed that the embedding dimensionality of hand kinematics did not exceed 8 even when humans individually moved the joints on their hands^77^. However, a more recent study showed evidence that even low-variance Principal Components contain some degree of task-relevant information, leading to the proposal that the distribution of eigenvalues of the covariance matrix of joint angles should not be truncated, and that the embedding dimensionality of everyday manual behaviors could be as high as 30^78^. Future studies should investigate how task complexity affects the dimensionality of the EMG signals associated with from very simple to highly dexterous hand gestures; such an investigation would help elucidate the current ambiguity in quantifying and interpreting behavioral complexity.

The use of linear models for decoding EMG signals might also be considered a limitation. Linear decoding is based on a weighted sum of neural signals; it is thus the projection of collective neural activity along a specific direction in the neural state space. Although this widely used approach^42,43,61^ has been superseded by nonlinear and recurrent alternatives^43,79–81^, linear decoders are simple, interpretable, and effective, and provide a useful tool for hypothesis testing. In this study, we were interested not in the absolute EMG decoding accuracy achieved when all neurons are used as inputs, but in the relative accuracy achieved when only the latent variables associated with the low-dimensional manifolds are used as inputs. Linear decoders were amply sufficient to quantify this comparison. Future studies, such as those involving neural prostheses, might benefit from the use of nonlinear and recurrent models of decoding for improved EMG decoding accuracy^81^, although the degree of improvement is likely to depend on task complexity and its effect on manifold nonlinearity.

## Conclusion

Our study demonstrates that the low dimensionality of task-specific manifolds in primary motor cortex is not limited to constrained laboratory tasks but is also present in unconstrained tasks. This finding suggests that the low dimensionality of M1 manifolds reflects an intrinsic computational property of M1, rather than being a byproduct of constrained movements. Even if the unconstrained tasks were relatively simple, to have established the existence of low-dimensional neural manifolds in M1 beyond our study.

Our study also revealed signatures of nonlinearity in the geometry of the M1 neural manifolds, indicating that the neural population activity can represent complex relationships between the latent variables. However, the population neural dynamics mostly explored nearly linear regions of the neural manifolds. This finding is consistent with our results demonstrating the ability to linearly decode EMG signals from relatively few linear latent variables, and provides a manifold-based understanding of the surprising effectiveness of linear methods in decoding the motor output in brain-machine interfaces^42,43,60,80^.

Additionally, our study emphasizes the importance of distinguishing between intrinsic and embedding dimensionalities when analyzing neural population activity. Our results suggest that the low embedding dimensionality of M1 manifolds reflects that the processing and communication requirements of M1 are relatively simple in comparison to those of other brain areas. For example, areas such as prefrontal cortex, parallel fibers in the cerebellum, and the primary visual cortex exhibit quite higher embedding dimensionalities. Future studies should explore the relationship between neural dimensionality, manifold nonlinearity, and information processing, both across different brain regions and across behavioral contexts of distinct complexity.

Overall, our study provides new insights into the computational properties of primary motor cortex and highlights the potential of low-dimensional representations for decoding motor output. The insights gained from this research have implications for extending the applicability of neural prosthetics and brain-machine interfaces to natural environments, while contributing to a broader understanding of how the motor cortex represents and processes movement-related information.

## REFERENCES

1. Gao, P. & Ganguli, S. On simplicity and complexity in the brave new world of large-scale neuroscience. Curr. Opin. Neurobiol. 32, 148–155 (2015).

2. Gallego, J. A., Perich, M. G., Miller, L. E. & Solla, S. A. Neural Manifolds for the Control of Movement. Neuron 94, 978–984 (2017).

3. Saxena, S. & Cunningham, J. P. Towards the neural population doctrine. Curr. Opin. Neurobiol. 55, 103–111 (2019).

4. Stopfer, M., Jayaraman, V. & Laurent, G. Intensity versus identity coding in an olfactory system. Neuron 39, 991–1004 (2003).

5. DiCarlo, J. J. & Cox, D. D. Untangling invariant object recognition. Trends Cogn. Sci. 11, 333–341 (2007).

6. Churchland, M. M. et al. Neural population dynamics during reaching. Nature 487, 51–56 (2012).

7. Mante, V., Sussillo, D., Shenoy, K. V. & Newsome, W. T. Context-dependent computation by recurrent dynamics in prefrontal cortex. Nature 503, 78–84 (2013).

8. Kaufman, M. T., Churchland, M. M., Ryu, S. I. & Shenoy, K. V. Cortical activity in the null space: permitting preparation without movement. Nat. Neurosci. 17, 440–448 (2014).

9. Sadtler, P. T. et al. Neural constraints on learning. Nature 512, 423–426 (2014).

10. Chung, S. Y., Lee, D. D. & Sompolinsky, H. Classification and geometry of general perceptual manifolds. Physical Review X (2018).

11. Remington, E. D., Narain, D., Hosseini, E. A. & Jazayeri, M. Flexible Sensorimotor Computations through Rapid Reconfiguration of Cortical Dynamics. Neuron 98, 1005–1019.e5 (2018).

12. Low, R. J., Lewallen, S., Aronov, D., Nevers, R. & Tank, D. W. Probing variability in a cognitive map using manifold inference from neural dynamics. bioRxiv 418939 (2018) doi:10.1101/418939.

13. Rubin, A. et al. Revealing neural correlates of behavior without behavioral measurements. Nat. Commun. 10, 4745 (2019).

14. Chaudhuri, R., Gerçek, B., Pandey, B., Peyrache, A. & Fiete, I. The intrinsic attractor manifold and population dynamics of a canonical cognitive circuit across waking and sleep. Nat. Neurosci. 22, 1512–1520 (2019).

15. Russo, A. A. et al. Neural Trajectories in the Supplementary Motor Area and Motor Cortex Exhibit Distinct Geometries, Compatible with Different Classes of Computation. Neuron 107, 745–758.e6 (2020).

16. Nieh, E. H. et al. Geometry of abstract learned knowledge in the hippocampus. Nature 595, 80–84 (2021).

17. Libby, A. & Buschman, T. J. Rotational dynamics reduce interference between sensory and memory representations. Nat. Neurosci. 24, 715–726 (2021).

18. Chandak, R. & Raman, B. Neural manifolds for odor-driven innate and acquired appetitive preferences. bioRxiv 2021.08.05.455310 (2021) doi:10.1101/2021.08.05.455310.

19. Ehrlich, D. B. & Murray, J. D. Geometry of neural computation unifies working memory and planning. Proc. Natl. Acad. Sci. U. S. A. 119, e2115610119 (2022).

20. Gardner, R. J. et al. Toroidal topology of population activity in grid cells. Nature 602, 123–128 (2022).

21. Churchland, M. M. & Shenoy, K. V. Temporal complexity and heterogeneity of single-neuron activity in premotor and motor cortex. J. Neurophysiol. 97, 4235–4257 (2007).

22. Vyas, S., Golub, M. D., Sussillo, D. & Shenoy, K. V. Computation Through Neural Population Dynamics. Annu. Rev. Neurosci. 43, 249–275 (2020).

23. Jazayeri, M. & Ostojic, S. Interpreting neural computations by examining intrinsic and embedding dimensionality of neural activity. Curr. Opin. Neurobiol. 70, 113–120 (2021).

24. Altan, E., Solla, S. A., Miller, L. E. & Perreault, E. J. Estimating the dimensionality of the manifold underlying multi-electrode neural recordings. PLoS Comput. Biol. 17, e1008591 (2021).

25. Gao, P. et al. A theory of multineuronal dimensionality, dynamics and measurement. bioRxiv 214262 (2017) doi:10.1101/214262.

26. Urai, A. E., Doiron, B., Leifer, A. M. & Churchland, A. K. Large-scale neural recordings call for new insights to link brain and behavior. Nat. Neurosci. 25, 11–19 (2022).

27. Perich, M. G., Gallego, J. A. & Miller, L. E. A Neural Population Mechanism for Rapid Learning. Neuron 100, 964–976.e7 (2018).

28. Gallego, J. A. et al. Cortical population activity within a preserved neural manifold underlies multiple motor behaviors. Nat. Commun. 9, 4233 (2018).

29. Campadelli, P., Casiraghi, E., Ceruti, C. & Rozza, A. Intrinsic Dimension Estimation: Relevant Techniques and a Benchmark Framework. Math. Probl. Eng. 2015, (2015).

30. Camastra, F. & Staiano, A. Intrinsic dimension estimation: Advances and open problems. Inf. Sci. 328, 26–41 (2016).

31. Horn, J. L. A Rationale and Test For The Number Of Factors In Factor Analysis. Psychometrika 30, 179–185 (1965).

32. Buja, A. & Eyuboglu, N. Remarks on Parallel Analysis. Multivariate Behav. Res. 27, 509–540 (1992).

33. Franklin, S. B., Gibson, D. J., Robertson, P. A., Pohlmann, J. T. & Fralish, J. S. Parallel Analysis: a method for determining significant principal components. J. Veg. Sci. 6, 99–106 (1995).

34. Facco, E., d’Errico, M., Rodriguez, A. & Laio, A. Estimating the intrinsic dimension of datasets by a minimal neighborhood information. Sci. Rep. 7, 12140 (2017).

35. Dinno, A. Exploring the Sensitivity of Horn’s Parallel Analysis to the Distributional Form of Random Data. Multivariate Behav. Res. 44, 362–388 (2009).

36. Levina, E. & Bickel, P. Maximum likelihood estimation of intrinsic dimension. Adv. Neural Inf. Process. Syst. 17, (2004).

37. Allegra, M., Facco, E., Denti, F., Laio, A. & Mira, A. Data segmentation based on the local intrinsic dimension. Sci. Rep. 10, 16449 (2020).

38. Denti, F., Doimo, D., Laio, A. & Mira, A. The generalized ratios intrinsic dimension estimator. Sci. Rep. 12, 20005 (2022).

39. Bac, J., Mirkes, E. M., Gorban, A. N., Tyukin, I. & Zinovyev, A. Scikit-Dimension: A Python Package for Intrinsic Dimension Estimation. Entropy 23, (2021).

40. Cunningham, J. P. & Yu, B. M. Dimensionality reduction for large-scale neural recordings. Nat. Neurosci. 17, 1500–1509 (2014).

41. Recanatesi, S. et al. Predictive learning as a network mechanism for extracting low-dimensional latent space representations. Nat. Commun. 12, 1417 (2021).

42. Carmena, J. M. et al. Learning to control a brain-machine interface for reaching and grasping by primates. PLoS Biol. 1, E42 (2003).

43. Glaser, J. I. et al. Machine Learning for Neural Decoding. eNeuro 7, (2020).

44. Faisal, A. A., Selen, L. P. J. & Wolpert, D. M. Noise in the nervous system. Nat. Rev. Neurosci. 9, 292–303 (2008).

45. Camastra, F. Data dimensionality estimation methods: a survey. Pattern Recognit. 36, 2945–2954 (2003).

46. Russo, A. A. et al. Motor Cortex Embeds Muscle-like Commands in an Untangled Population Response. Neuron 97, 953–966.e8 (2018).

47. Pashkovski, S. L. et al. Structure and flexibility in cortical representations of odour space. Nature 583, 253–258 (2020).

48. Stringer, C., Michaelos, M., Tsyboulski, D., Lindo, S. E. & Pachitariu, M. High-precision coding in visual cortex. Cell 184, 2767–2778.e15 (2021).

49. Churchland, M. M., Cunningham, J. P., Kaufman, M. T., Ryu, S. I. & Shenoy, K. V. Cortical preparatory activity: representation of movement or first cog in a dynamical machine? Neuron 68, 387–400 (2010).

50. Kato, S. et al. Global brain dynamics embed the motor command sequence of Caenorhabditis elegans. Cell 163, 656–669 (2015).

51. Zimnik, A. J. & Churchland, M. M. Independent generation of sequence elements by motor cortex. Nat. Neurosci. 24, 412–424 (2021).

52. Fusi, S., Miller, E. K. & Rigotti, M. Why neurons mix: high dimensionality for higher cognition. Curr. Opin. Neurobiol. 37, 66–74 (2016).

53. Badre, D., Bhandari, A., Keglovits, H. & Kikumoto, A. The dimensionality of neural representations for control. Curr Opin Behav Sci 38, 20–28 (2021).

54. Maass, W. Searching for principles of brain computation. Current Opinion in Behavioral Sciences 11, 81–92 (2016).

55. Semedo, J. D., Zandvakili, A., Machens, C. K., Yu, B. M. & Kohn, A. Cortical Areas Interact through a Communication Subspace. Neuron 102, 249–259.e4 (2019).

56. Kohn, A. et al. Principles of Corticocortical Communication: Proposed Schemes and Design Considerations. Trends Neurosci. 43, 725–737 (2020).

57. Rigotti, M. et al. The importance of mixed selectivity in complex cognitive tasks. Nature 497, 585–590 (2013).

58. Kaufman, M. T. et al. The implications of categorical and category-free mixed selectivity on representational geometries. Curr. Opin. Neurobiol. 77, 102644 (2022).

59. Stringer, C., Pachitariu, M., Steinmetz, N., Carandini, M. & Harris, K. D. High-dimensional geometry of population responses in visual cortex. Nature 571, 361–365 (2019).

60. Pandarinath, C. et al. Inferring single-trial neural population dynamics using sequential auto-encoders. Nat. Methods 15, 805–815 (2018).

61. Pandarinath, C. et al. Latent Factors and Dynamics in Motor Cortex and Their Application to Brain-Machine Interfaces. J. Neurosci. 38, 9390–9401 (2018).

62. Lanore, F., Cayco-Gajic, N. A., Gurnani, H., Coyle, D. & Silver, R. A. Cerebellar granule cell axons support high-dimensional representations. Nat. Neurosci. 24, 1142–1150 (2021).

63. Musslick, S., et al. Multitasking capability versus learning efficiency in neural network architectures. in (Cognitive Science Society, 2017).

64. Musslick, S. & Cohen, J. D. A mechanistic account of constraints on control-dependent processing: Shared representation, conflict and persistence. https://cogsci.mindmodeling.org/2019/papers/0161/0161.pdf (2019).

65. Pryluk, R., Kfir, Y., Gelbard-Sagiv, H., Fried, I. & Paz, R. A Tradeoff in the Neural Code across Regions and Species. Cell 176, 597–609.e18 (2019).

66. Sagiv, Y., Musslick, S., Niv, Y. & Cohen, J. D. Efficiency of learning vs. processing: Towards a normative theory of multitasking. arXiv [q-bio.NC] (2020).

67. Boser, B. E., Guyon, I. M. & Vapnik, V. N. A training algorithm for optimal margin classifiers. in Proceedings of the fifth annual workshop on Computational learning theory 144–152 (Association for Computing Machinery, 1992).

68. Cohen, U., Chung, S., Lee, D. D. & Sompolinsky, H. Separability and geometry of object manifolds in deep neural networks. Nat. Commun. 11, 746 (2020).

69. Litwin-Kumar, A., Harris, K. D., Axel, R., Sompolinsky, H. & Abbott, L. F. Optimal Degrees of Synaptic Connectivity. Neuron 93, 1153–1164.e7 (2017).

70. Mastrogiuseppe, F. & Ostojic, S. Linking Connectivity, Dynamics, and Computations in Low-Rank Recurrent Neural Networks. Neuron 99, 609–623.e29 (2018).

71. Schuessler, F., Dubreuil, A. & Mastrogiuseppe, F. Dynamics of random recurrent networks with correlated low-rank structure. Physical Review (2020).

72. Beiran, M., Dubreuil, A., Valente, A., Mastrogiuseppe, F. & Ostojic, S. Shaping Dynamics With Multiple Populations in Low-Rank Recurrent Networks. Neural Comput. 33, 1572–1615 (2021).

73. Pollock, E. & Jazayeri, M. Engineering recurrent neural networks from task-relevant manifolds and dynamics. PLoS Comput. Biol. 16, e1008128 (2020).

74. Langdon, C., Genkin, M. & Engel, T. A. A unifying perspective on neural manifolds and circuits for cognition. Nat. Rev. Neurosci. (2023) doi:10.1038/s41583-023-00693-x.

75. Bhandari, A., Gagne, C. & Badre, D. Just above Chance: Is It Harder to Decode Information from Prefrontal Cortex Hemodynamic Activity Patterns? J. Cogn. Neurosci. 30, 1473–1498 (2018).

76. Bialek, W. What do we mean by the dimensionality of behavior? arXiv preprint arXiv:2008.09574 (2020).

77. Todorov, E. & Ghahramani, Z. Analysis of the synergies underlying complex hand manipulation. Conf. Proc. IEEE Eng. Med. Biol. Soc. 2004, 4637–4640 (2004).

78. Yan, Y., Goodman, J. M., Moore, D. D., Solla, S. A. & Bensmaia, S. J. Unexpected complexity of everyday manual behaviors. Nat. Commun. 11, 3564 (2020).

79. Glaser, J. I., Benjamin, A. S., Farhoodi, R. & Kording, K. P. The roles of supervised machine learning in systems neuroscience. Prog. Neurobiol. 175, 126–137 (2019).

80. Perkins, S. M., Cunningham, J. P., Wang, Q. & Churchland, M. M. Simple decoding of behavior from a complicated neural manifold. bioRxiv 2023.04.05.535396 (2023) doi:10.1101/2023.04.05.535396.

81. Deo, D. R., et al. Translating deep learning to neuroprosthetic control. bioRxiv (2023) doi:10.1101/2023.04.21.537581.

