## Supplementary Figures for "Low-dimensional neural manifolds for the control of constrained and unconstrained movements"

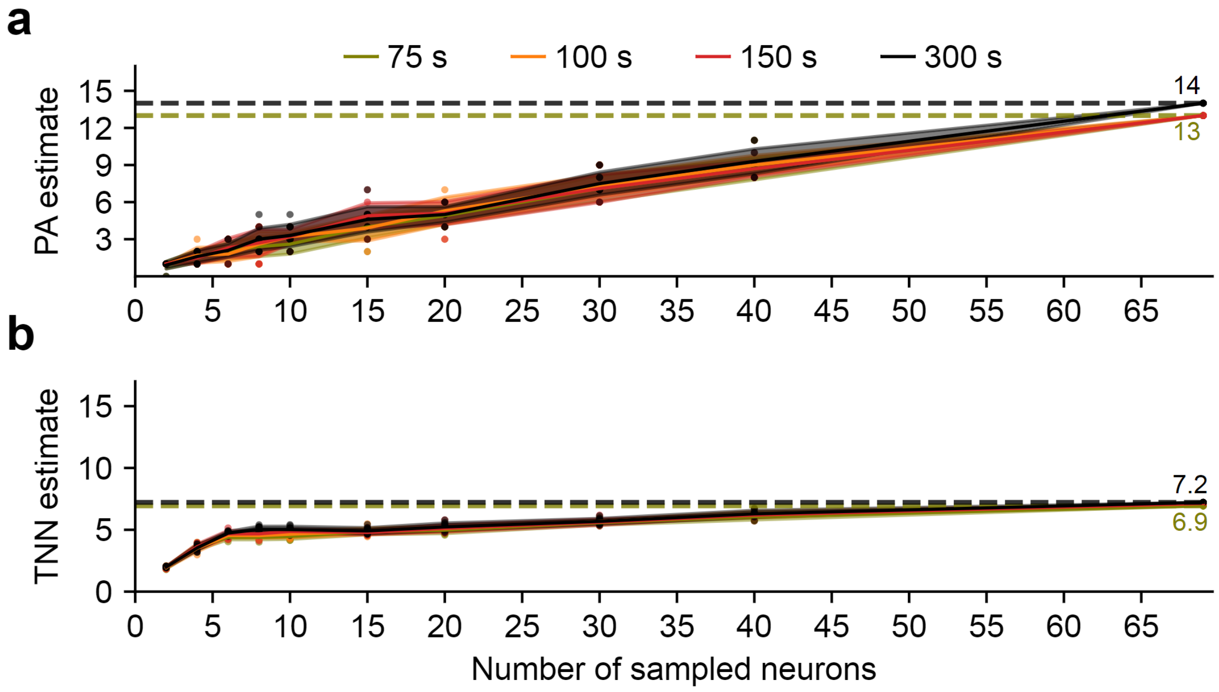


**Supplementary Figure 1:** The effect of temporal length of data and number of neurons on estimating embedding and intrinsic dimensionalities. **a**) Parallel Analysis (PA) estimates for the embedding dimensionality with increasing lengths of temporal data and number of neurons for Monkey P doing the bar walk. Colors represent the different amounts of temporal data used for the dimensionality estimate, ranging from 1500 bins of neural data corresponding to 75 seconds (binned at 50 ms) to 6000 bins of neural data corresponding to 300 seconds. The analysis was repeated 10 times, sampling a subset of neurons ranging from 2 to 69 (maximum neurons available for this session). b) Same as in panel a, but Two Nearest-Neighbors (TNN) estimates for the intrinsic dimensionality instead of PA estimates for the embedding dimensionality.

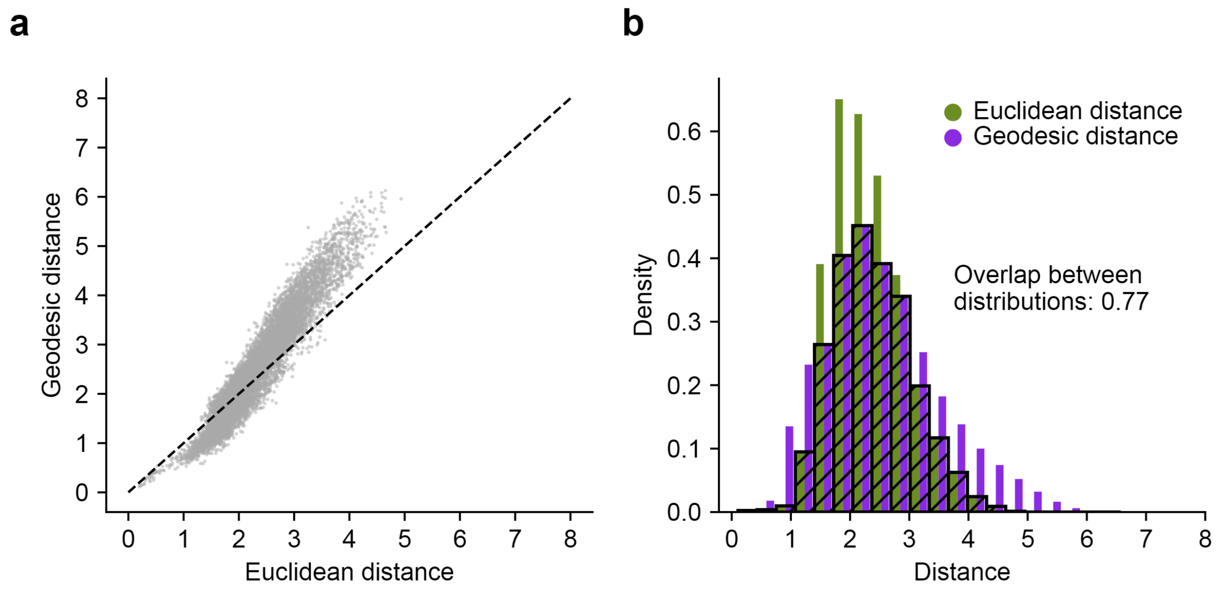


**Supplementary Figure 2**: Computation of the local flatness index. a) Comparison between the Euclidean and geodesic distances for all pairs of neural population data points recorded for Monkey P while performing the bar walk task. b) The distributions of Euclidean (olive) and geodesic (violet) distances for the same task. The distributions were normalized to represent probability densities. The sum of the overlap between the two distributions (shown as hatched black bars) is the local flatness index, equal to 0.77 for this dataset.

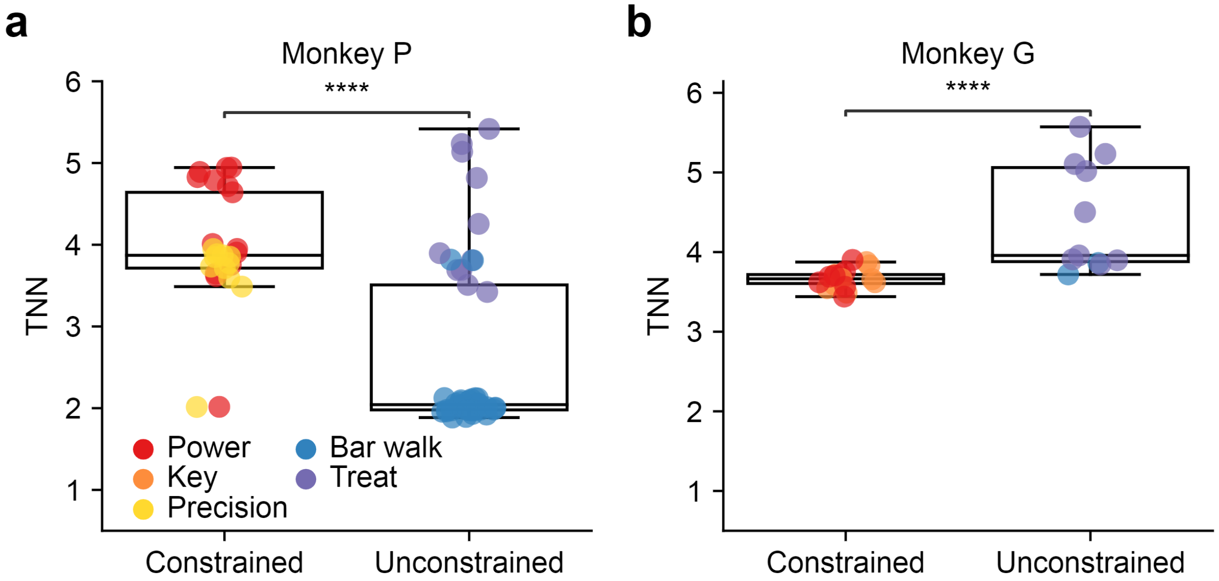


**Supplementary Figure 3:** Intrinsic dimensionality of EMG signals. The intrinsic dimensionality of the EMG signals was estimated using Two Nearest-Neighbors (TNN). Different colors represent different tasks. Warm colors indicate data collected in the constrained laboratory setting and cold colors indicate data collected in the unconstrained cage setting. a) Intrinsic dimensionality of EMG for Monkey P ($p \approx 0$). b) Same as in panel a, but for Monkey G ($p \approx 0$).

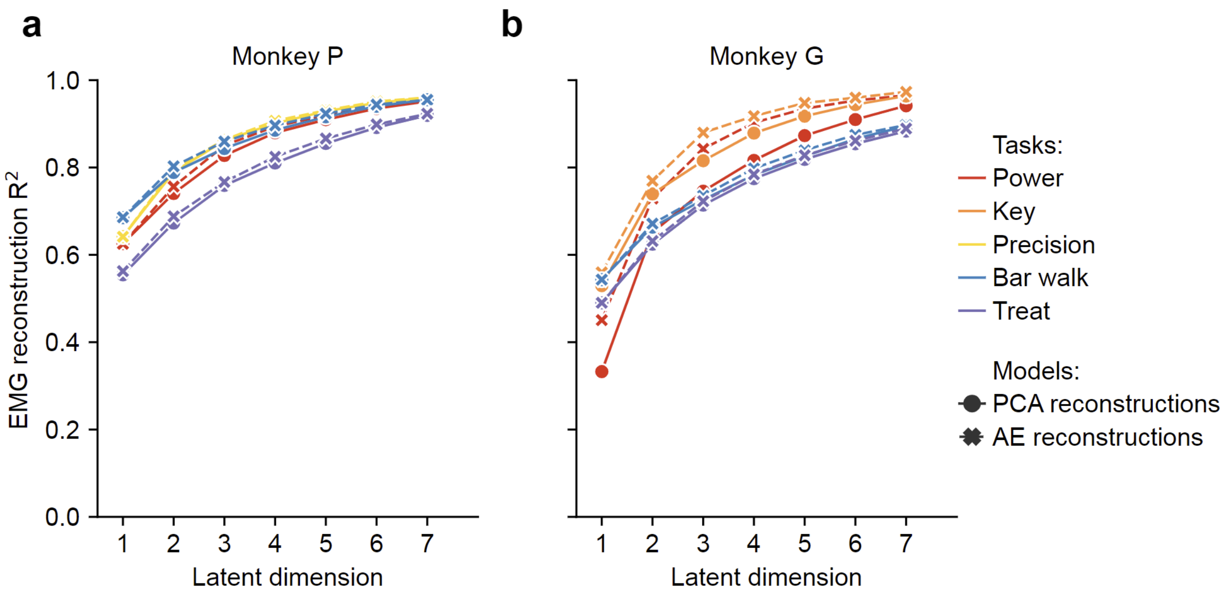


**Supplementary Figure 4:** EMG reconstruction accuracies with progressively increasing EMG manifold dimensionality. a) We progressively increased the latent dimensionality of the linear (Principal Component Analysis, PCA) and nonlinear (autoencoder, AE) EMG manifolds of Monkey P performing a variety of tasks. Each color corresponds to a different task. All results for a given task were averaged. Circle and cross symbols indicate the reconstruction accuracy of the EMG data from the low dimensional EMG manifolds of varying latent dimensionality using PCA and autoencoder (AE), respectively. b) Same as in panel a but for Monkey G.

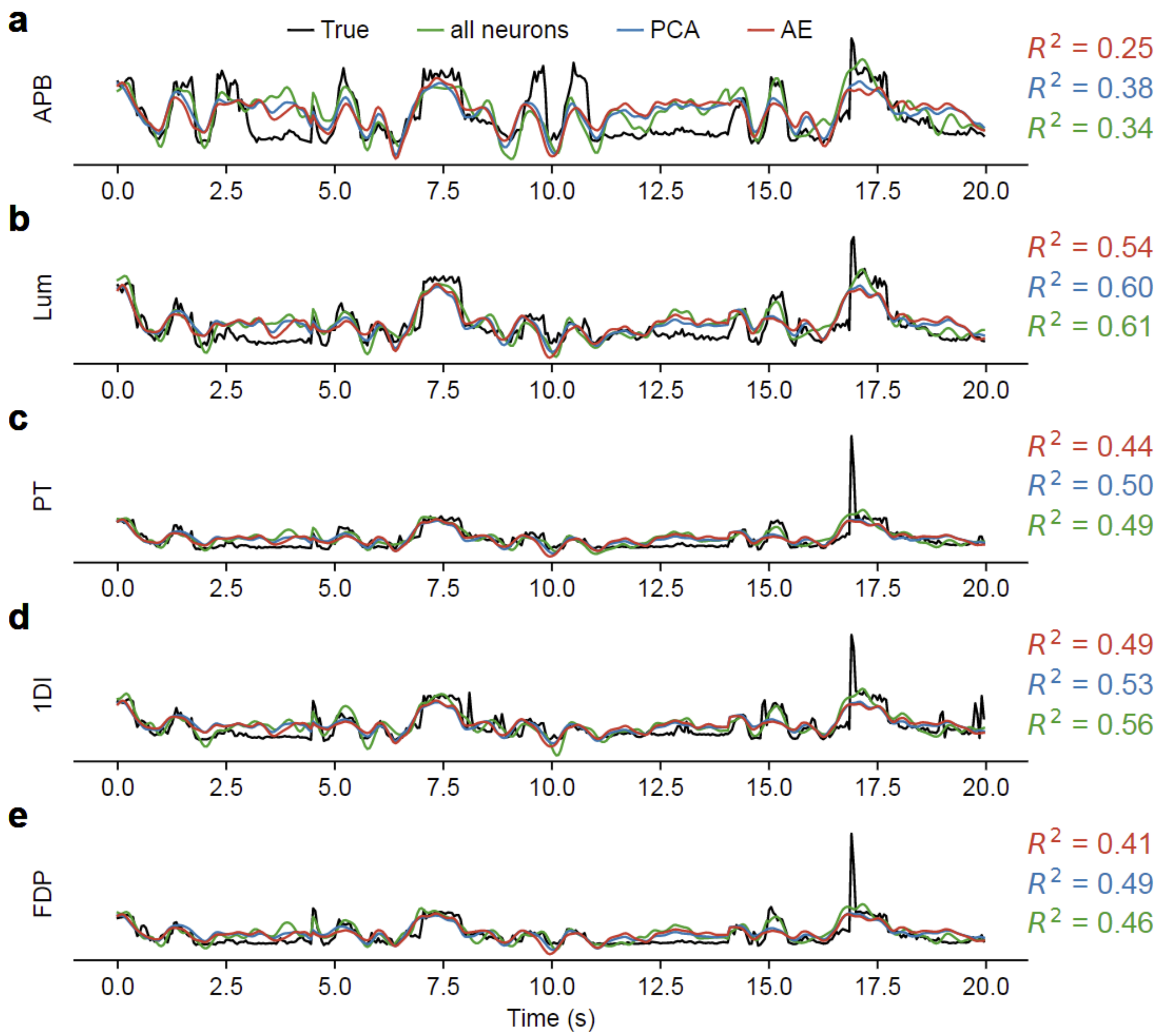


**Supplementary Figure 5:** EMG prediction examples for Monkey P performing the bar walk task in the cage. Each panel corresponds to the predictions of intramuscular EMG recorded from a different muscle. Black traces represent the normalized EMG activity. Green traces represent EMG activity predictions using all recorded neurons. Blue and red traces represent EMG predictions from low-dimensional linear (PCA) and nonlinear (autoencoder) latent variables, respectively. Corresponding $R^2$ values are denoted on the right of each panel. EMG predictions for a) Abductor Pollicis Brevis (APB), b) Lumbricals (Lum), c) Pronator teres (PT), d) First dorsal interossei (1DI), e) Flexor Digitorum Profundus (FDP).

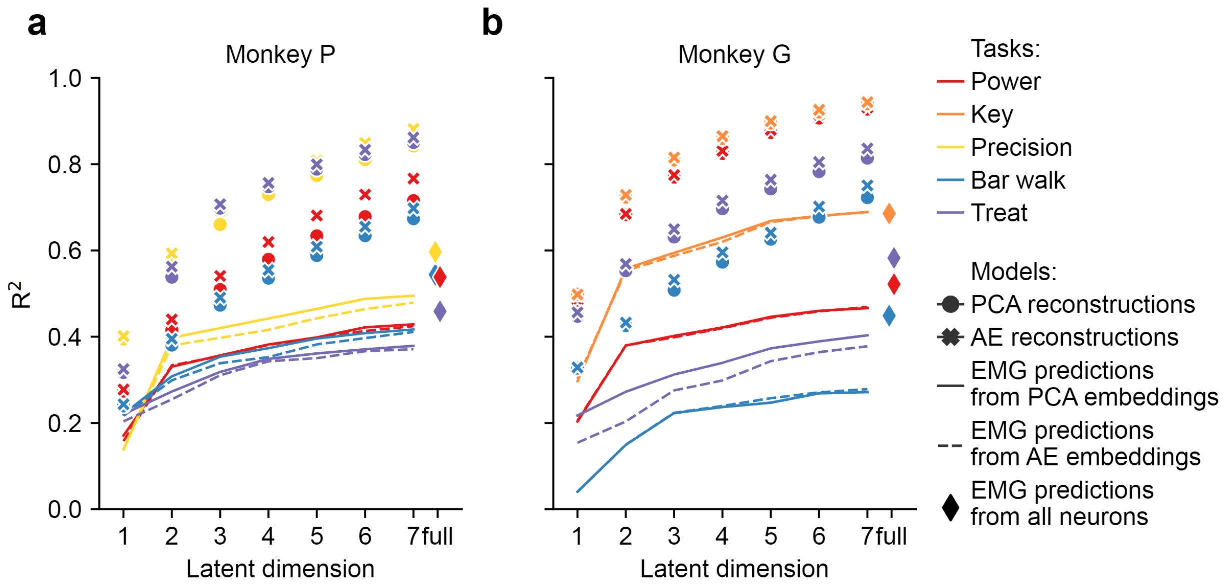


**Supplementary Figure 6:** EMG predictions and neural reconstruction accuracies with progressively increasing neural manifold dimensionality. a) We progressively increased the latent dimensionality of the linear (Principal Component Analysis, PCA) and nonlinear (autoencoder, AE) neural manifolds of Monkey P performing a variety of tasks. Each color corresponds to a different task. All results for a given task were averaged. Solid lines indicate average EMG predictions from linear neural manifolds of varying latent dimensionality. Dashed lines indicate average EMG predictions from nonlinear neural manifolds of varying latent dimensionality. Diamond symbols on the right indicate the average EMG predictions using all available neurons. All EMG predictions shown in the figure are the averages of the test folds of five-fold cross validation. Circle and cross symbols indicate the reconstruction accuracy of the neural data from the low-dimensional neural manifolds of varying latent dimensionality using PCA and autoencoder (AE), respectively. b) Same as in panel a but for Monkey G.
